# Deciphering enhancer sequence using thermodynamics-based models and convolutional neural networks

**DOI:** 10.1101/2021.03.01.433444

**Authors:** Payam Dibaeinia, Saurabh Sinha

## Abstract

Deciphering the sequence-function relationship encoded in enhancers holds the key to interpreting non-coding variants and understanding mechanisms of transcriptomic variation. Several quantitative models exist for predicting enhancer function and underlying mechanisms; however, there has been no systematic comparison of these models characterizing their relative strengths and shortcomings. Here, we interrogated a rich data set of neuroectodermal enhancers in *Drosophila*, representing cis- and trans- sources of expression variation, with a suite of biophysical and machine learning models. We performed rigorous comparisons of thermodynamics-based models implementing different mechanisms of activation, repression, and cooperativity. Moreover, we developed a convolutional neural network (CNN) model, called CoNSEPT, that learns enhancer “grammar” in an unbiased manner. CoNSEPT is the first general-purpose CNN tool for predicting enhancer function in varying conditions, and we show that such complex models can suggest interpretable mechanisms. We found model-based evidence for mechanisms previously established for the studied system, including cooperative activation and short-range repression. The data also favored one hypothesized activation mechanism over another and suggested an intriguing role for a direct, distance-independent repression mechanism. Our modeling shows that while fundamentally different models can yield similar fits to data, they vary in their utility for mechanistic inference. CoNSEPT is freely available at: https://github.com/PayamDiba/CoNSEPT.

## Introduction

Transcriptional regulation in metazoans is mediated by proteins called transcription factors (TF) that bind to specific sites in regulatory regions called enhancers (1), via TF-DNA interactions and cooperative DNA binding (2). Many TFs that occupy their respective binding sites interact with each other and with the transcription start site over long and short distances to influence the recruitment of transcription machinery and transcription initiation (3). These simultaneous interactions establish a complex regulatory code that drives a gene’s expression in varying cellular conditions or cell types.

Gene regulatory mechanisms encoded in an enhancer can be fairly complex and have been the subject of numerous studies, notably the detailed experimental dissection of developmental enhancers in Drosophila (4–6). Such studies have significantly advanced our understanding of regulatory mechanisms. For example, certain TFs that are known to inhibit transcription (“repressors”) have been found to function only if bound at short distances from activator binding sites (7–9). As another example, TFs that are responsible for promoting transcription (“activators”) have been shown in some cases to contribute synergistically to the gene’s expression (10, 11), possibly due to cooperative DNA-binding by TFs to adjacent binding sites. These complex regulatory mechanisms may be mirrored in rules underlying the arrangement of binding sites in an enhancer, a phenomenon sometimes called cis-regulatory “grammar” (12–15). Precise characterization of the sequence-function relationship encoded in enhancers therefore requires interpreting how a collection of binding sites for one or more TFs works together and how such combinatorial action is influenced by site arrangements as well as varying TF concentrations in different cellular contexts. The challenge goes beyond a general understanding of the underlying principles (e.g., “TF X is a short-range repressor” or “TFs X and Y activate synergistically”): often, one seeks a quantitative model capable of predicting a given enhancer’s regulatory output in varying cellular conditions and the effect of sequence variations such as disease-related non-coding polymorphisms within the enhancer (16). Indeed, such “sequence-to-expression models” are an active area of research today (17).

The most direct efforts to deciphering the cis-regulatory code of enhancers have been through experiments that record expression readouts of many variants of an enhancer (14, 18) or many enhancers under similar control (12, 19, 20). Often, the variants are synthetic constructs that manifest a diversity of TF binding site composition and arrangement; in some cases, their functions (expression level) are determined in a single cellular context (12) while occasionally the expression readout is obtained in varying cellular contexts, e.g., nuclei of the early *Drosophila* embryo (5, 20–22) or *Ciona* embryos. Such data sets are then analyzed through a specialized mathematical model whose structure incorporates mechanistic hypotheses and whose parameters represent quantitative details of those mechanisms, such as activation strength and distance-dependence of cooperative and repressive interactions among binding sites (17, 18, 21, 23–27). The most effective frameworks for such mathematical mechanistic modeling have been based on equilibrium thermodynamics (14, 28, 29), although non-equilibrium models have also been motivated and proposed (30–32).

Different thermodynamics-based models implement regulatory mechanisms in different ways. For example, some models accord the activation by a TF exclusively to its DNA-binding ability (18, 25), while others (24, 28) model activation by additional free parameters (beyond those for TF-DNA binding) that explicitly represent long-range interactions between activators and the basal transcription machinery (BTM). Similarly, different models employ various representations of short-range repression, e.g., via quenching of a bound activator or mediating local chromatin remodeling through recruitment of co-repressors (18, 21, 25, 28), and some even accommodate longer-range repression via direct inhibitory interaction between the bound repressor and BTM (28, 33). Typically, an enhancer dataset is analyzed using a mathematical model with a pre-determined structure (qualitative mechanisms) and its parameters are tuned to fit the data. It is rare for the same study to investigate models with varying assumptions about regulatory mechanisms to determine those best supported by the data. Here, we sought to bridge this gap by asking if various existing formalisms differ in their ability to model sequence-to-expression relationships.

In addition to thermodynamics-based models, statistical and machine learning (ML) models have also provided useful quantitative descriptions of cis-regulatory encoding (34, 35). Such models typically avoid strong pre-conceptions about underlying mechanisms, providing a “data-driven” approach to quantitative modeling of enhancers, as a counterpoint to the “hypothesis-driven” approach of thermodynamics-based models. Recently, Avsec et al. (36) trained a deep neural network model on TF-DNA binding data and showed how interrogating the trained model can reveal mechanistic insights. While neural network models have been frequently applied to TF-DNA binding and epigenomic data (37–39), their utility for sequence-to-expression modeling remains to be demonstrated. This is partly because data sets with direct expression measurements on enhancers (14, 18, 28) in diverse conditions are of relatively modest sizes and the models tend to be heavily parameterized. Thus, the second major motivation of this study was to test if ML models and especially neural network models provide a practical alternative to thermodynamics-based models for enhancer sequence-function relationships, and to assess their relative merits and weaknesses.

Motivated by the above considerations, we tested a suite of quantitative models, including linear models, thermodynamics-based models, and a newly developed convolutional neural network (CNN) model, on a rich sequence-expression data set previously reported by Sayal et al. (14). The data include expression measurements of an enhancer that drives expression of the *rhomboid (rho)* gene in the *D. melanogaster* embryo, along with several synthetic variants of the enhancer and several endogenous enhancers with similar regulatory function. Importantly, the dataset not only represents variation of enhancer sequences, it also includes changes in cellular conditions.

We used rigorous model comparisons and prior mechanistic studies of this regulatory system to evaluate the suite of quantitative models. The thermodynamics-based models we tested included two different activation mechanisms, three repression mechanisms, and the presence or absence of cooperative activation via proximal binding sites. We tested linear and generalized linear models that combine binding site contributions additively, as well as an extension where pairwise TF interactions were allowed. Finally, we developed and tested a new neural network model for the expression driven by a given enhancer sequence in varying cellular conditions. This model, called “CoNSEPT” (**Co**nvolutional **N**eural Network-based **S**equence-to-**E**xpression **P**rediction **T**ool), utilizes user-provided DNA binding motifs and condition- or cell type-specific concentrations of TFs, and can thus quantify the regulatory role of each TF. Importantly, it learns salient rules of binding site arrangement in a purely data-driven manner, without presuming any particular distance-dependence function of TF-TF interaction as is commonly done in thermodynamics-based models.

Our modeling shows that the three broad categories of models are competitive with each other in terms of their ability to fit the enhancer-expression data set. We found that convolutional neural networks can be reliably trained on relatively modest-sized training data and can learn aspects of cis-regulatory grammar in a fully data-driven manner. To our knowledge, this is the first demonstration of a CNN predicting expression variation across sequences as well as across conditions. Our thermodynamics-based modeling showed that explicitly modeling the strength of activators is advantageous compared to ascribing activator strength solely to its DNA-binding, but that different biochemical mechanisms of short-range repression cannot be reliably distinguished based on the dataset. Both thermodynamics-based and neural network models detected a significant role for cooperative interaction between activator sites. Intriguingly, both types of models suggested a potential role for a direct repression mechanism that is not short-range, the predominant theory of repressor action for this system. Our baseline linear models showed good agreement with data but were, by design, limited in offering mechanistic insights into the cis-regulatory code, including rules of binding site arrangements and interactions. Overall, this work conveys a positive outlook for the modeler, who has at their disposal a variety of tools of varying complexity with which to understand a regulatory system at the level of enhancer sequences.

## Results

### A gene expression data set with cis and trans variations

We analyzed a data set generated and first modeled by Sayal et. al. (14). It includes expression levels driven by a well-studied enhancer of the gene *rhomboid (rho)* (**Figure 1A**, henceforth called the *rho* enhancer) in the early *D. melanogaster* embryo. This enhancer is regulated by three transcription factors (TFs): Dorsal (DL), Twist (TWI) and Snail (SNA). DL and TWI are known to activate and SNA represses *rho* expression (4, 14). Binding sites of these TFs in the *rho* enhancer are well mapped and shown in Figure 1A. The expression levels of *rho* driven by the wild-type (WT) enhancer were measured by Sayal et al. as a function of cellular (nuclear) position along the ventral-dorsal (V-D) axis of the embryo (**Figure 1B**), using reporter assays. The enhancer’s expression profile is quantitatively represented by a 17-dimensional vector, where the 17 dimensions represent uniformly spaced positions or “bins” along the V-D axis from the ventral end to 40% of the V-D axis length. Concentration profiles of the three TFs are also available, as 17-dimensional vectors analogous to the enhancer expression profiles (Figure 1B). Moreover, similar expression profiles were generated for 37 synthetic variants of the WT *rho* enhancer, where each variant was constructed by mutagenesis of one or multiple TF binding sites (**Figure 1C**). (Our nomenclature for the enhancers is different from that of Sayal et al., see Supplementary Table S1.) Thus, the data set captures expression variation across different trans contexts (cells at different V-D axis positions, with varying TF levels) as well as different cis contexts (WT enhancer and synthetic variants).

**Figure 1:**
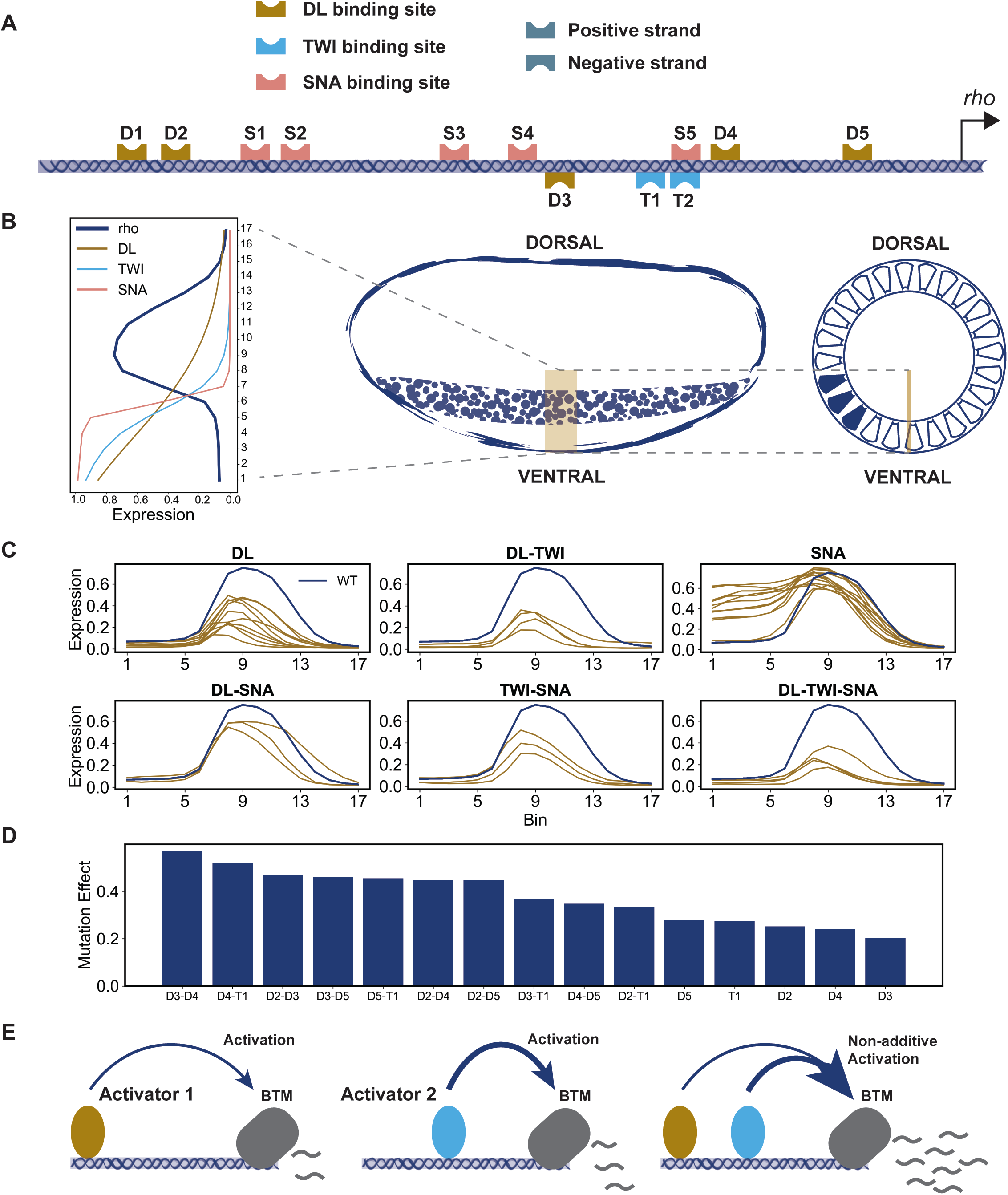
Overview of the data used in this study. (A) Schematic of the wild-type (WT) enhancer of the *Drosophila rhomboid* (*rho*) gene. Binding sites of the three TFs were identified using the PWMs employed in (14). All annotated sites agree with those found in (14) and except for D1 and S2 all the sites match with the in vitro footprinted sites characterized previously (14). (B) The levels of the three regulators and the expression of *rho* driven by the wild-type enhancer in 17 equidistant points along 0-40% of ventral-dorsal (V-D) axis. (C) The expression of *rho* driven by perturbed enhancers (shown in brown) representing mutagenesis of binding sites of one or more TFs. (D) Each activator’s site deletion (or combination thereof) is expected to reduce peak expression of rho (at bin 8 on the V-D axis); we therefore defined the effect of a variant enhancer (Y-axis) as the difference between the expression driven by it and the wild-type expression at this position of the axis. The effect of T2 single site deletion is not shown due to its overlap with the SNA site S5. (E) Schematic of synergistic activation, where the activation driven by two bound activators (right) is greater than the sum of their individual activation effects (left and middle).

**Figure 1D** reports on a selection of the synthetic enhancers – those representing DL and/or TWI site deletions, showing the difference in activation by each enhancer compared to the WT enhancer. Interestingly, deleting the two sites with smallest individual effects (“D3” and “D4”) simultaneously has the largest effect among all variants. Two other sites – “D2” and “T1” – individually have at least as much contribution as D3 and D4 individually, but their simultaneous deletion has a substantially smaller effect than the D3-D4 double deletion. This and other aspects of Figure 1D suggest non-linear regulatory contributions (**Figure 1E**) from sites in the *rho* enhancer, and present an interesting challenge for current mathematical models of cis-regulatory encoding: *can existing models capture the subtle variations of function encoded in these variant enhancers, and if so, can they reveal new insights about the underlying regulatory mechanisms?*

To meet the above challenge, we trained diverse mathematical models that map TF concentration profiles and enhancer sequence to the enhancer’s expression profile, for all 38 enhancers (WT and 37 variants, henceforth called the “training set”) simultaneously. The accuracy, or “goodness-of-fit”, of a model was measured by the root mean squared error (RMSE) between predicted and real expression profiles along the V-D axis, averaged over all enhancers; this is referred to as the “train error” below. The data set also includes expression profiles of 13 other enhancers that have expression profiles similar to *rho* (Supplementary Figure S1); these are orthologs of the *rho* enhancer from other Drosophila species or enhancers of other *D. melanogaster* genes with a neuroectodermal expression pattern similar to *rho*. We used these 13 enhancers, which represent greater sequence diversity than do the 38 enhancers in the “training set” (above), as the “test set”. The average RMSE of a model on these enhancers is referred to as “test error” below. By fitting, evaluating and comparing various models that differ in their explicit encoding of biophysical mechanisms, we hoped to draw inferences about specific mechanisms that are supported by the data set.

### Linear models provide a good phenomenological baseline

We first trained a linear model (LM) to assess the baseline explanatory power of statistical models on the Sayal et al. data set. In these models (35), expression driven by an enhancer is the sum of contributions from all TFs (**Figure 2A**). The contribution of a TF is the product of the TF’s concentration, its binding site strength at the enhancer estimated using its Position Weight Matrix (PWM) (see Methods) and a tunable parameter that represents the TF’s regulatory strength and direction (activator/repressor). We also tested a generalized linear model (GLM) (23, 40), where expression is a sigmoidal function of the sum of TF contributions (see Methods), such that the response of a gene is less sensitive to the concentration of its regulators at very low or high concentrations. Both LM and GLM have only one free parameter per TF and are the simplest of the models evaluated here. To impose our prior knowledge of TF roles, the trainable weights for DL and TWI were restricted to positive values and the weight for SNA was restricted to negative values. For LM we computed the globally optimal parameters (on the training set) while for GLM an ensemble of 100 models was trained.

**Figure 2:**
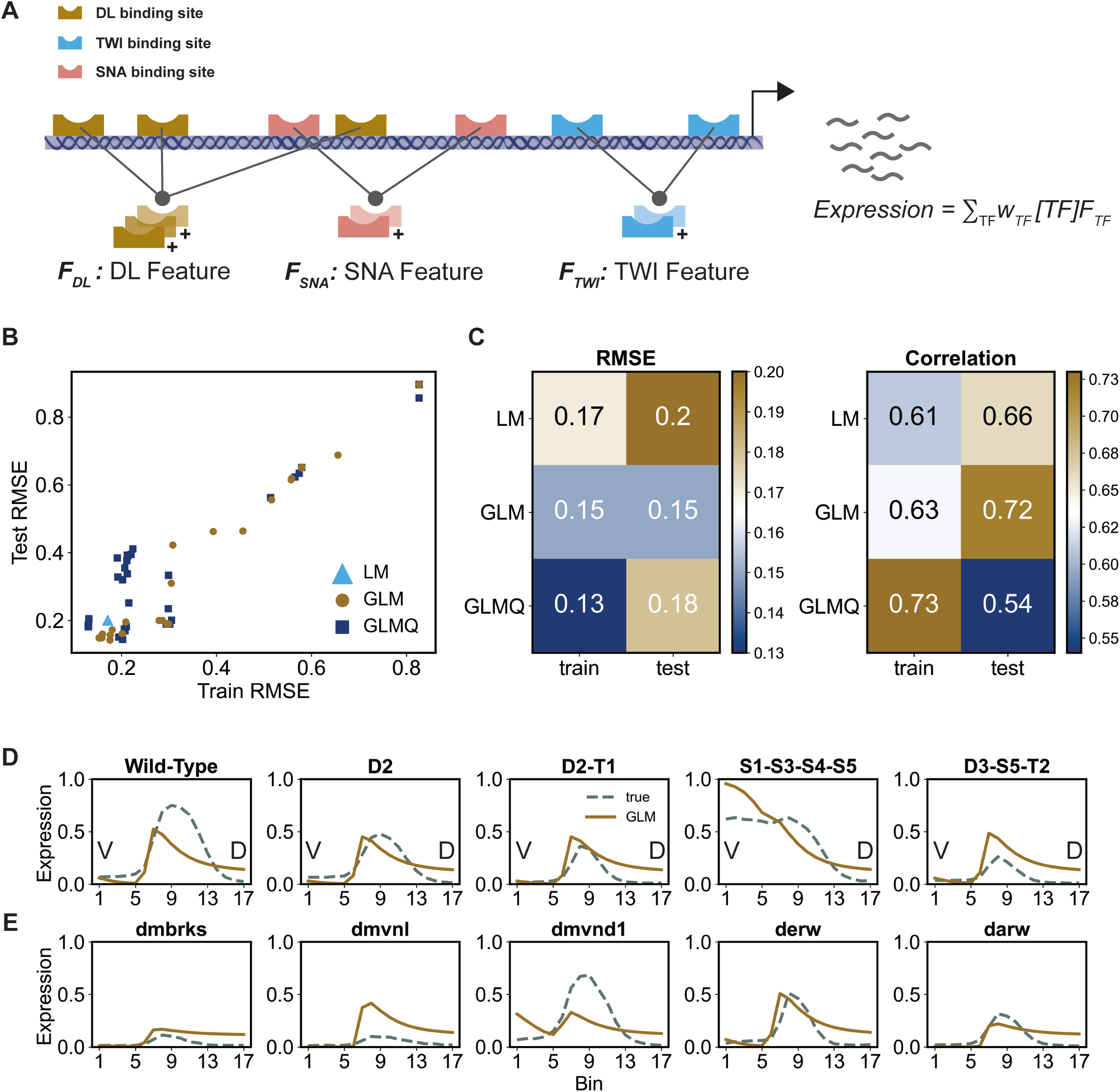
Linear and generalized linear models of gene expression. (A) Schematic representation of linear model (LM). Each TF’s “feature” is obtained by aggregating the strengths of all of its binding sites in the enhancer. Expression is modeled as the weighted (*w*) sum of all TF features multiplied by their corresponding concentration profile ([.]). (B) Train and Test RMSE for LM and the ensemble of GLM (generalized linear model) and GLMQ (GLM with quadratic terms) models. (C) RMSE and correlation scores of LM and best-fit (smallest train RMSE) GLM and GLMQ models in train and test data. (D) Predictions of best-fit GLM model on select train enhancers (brown curves) shown versus true expressions (blue curves). Left-most panel corresponds to the Wild-Type enhancer and the remaining panels correspond to perturbed enhancers titled by the site deletion they represent (e.g., “D2-T1” corresponds to simultaneous deletion of DL site D2 and TWI site T1). (E) Predictions of best-fit GLM model on select test enhancers (brown curves) shown versus true expressions (blue curves).

**Figure 2B** shows the train and test errors (RMSE) and correlation coefficient between real and predicted expression profiles, for LM as well as the ensemble of GLM models. The GLM model with the smallest train error was selected from the ensemble and its performance was compared against the LM model (**Figure 2C**). The GLM model clearly shows better fits than LM model in terms of error (RMSE) and correlation on both train and test data sets. Examining its predictions more closely (**Figure 2D, E**), we find that the GLM model often correctly predicts the main features of an enhancer’s readout, e.g., location of expression peak along the axis, but is also prone to predicting excessive expression at the dorsal end, which is due to inaccurate estimation of the basal transcription level (the intercept term in the linear function).

The above models consider each enhancer as a “bag of sites” (41) where multiple TFs and TF sites contribute additively to the regulatory output. In an attempt to capture any non-additive contributions from pairs of TFs, as is believed to arise from cooperative activity (11), we extended the GLM model (‘GLMQ’) to include quadratic terms that represent products of TF concentrations (see Methods). Though the training RMSE improved compared to GLM due to the additional parameters, the test RMSE and test correlation deteriorated substantially (**Figure 2C**). Notably, GLMQ shows a negative trained coefficient corresponding to DL-TWI interaction (see Supplementary table S2 for trained parameters) which contradicts the previous findings of the synergistic activity between these two activators reported in literature (42). In summary, cooperative interactions were not reliably learnt by simply adding quadratic terms to the generalized linear model. This is not surprising, since this approach to modeling TF cooperativity does not consider dependence of the interactions on inter-site distances (2).

### Thermodynamics-based models reveal biochemical mechanisms

We next employed thermodynamics-based models of gene expression (14, 28) that explicitly incorporate biochemical mechanisms of gene regulation, including TF-DNA binding affinities, activation and repression mechanisms, synergistic activity of multiple TFs, and distance-dependent interactions between TFs bound at proximal sites. Different models, representing different combinations of mechanisms, were implemented as variations of the same thermodynamics-based sequence-to-expression modeling framework, called GEMSTAT (28). Evaluation of different GEMSTAT model variations was performed with the same parameter fitting techniques, and the compared models typically shared many parameters, differing only in the desired mechanistic aspect, making their comparison more controlled.

#### Activation mechanisms

We first examined and compared two different activation mechanisms that have been implemented in past modeling studies. In both mechanisms, activation is due to binding of activator TFs to the enhancer and stronger binding leads to greater activation. In one class of models, e.g., that used by Sayal et al. in their original analysis of the data set (14), the contribution of an activator binding site depends only on the binding affinity of the site for its cognate TF and the TF’s concentration. (For now, we ignore effects of other binding sites on this site’s contribution.) These two factors together determine the fractional “occupancy” of the site by the TF, and its contribution to expression depends solely on its occupancy. In the second class of models, the site’s contribution additionally depends on the particular TF’s “potency”, which may be different for different TFs. That is, two sites with the same fractional occupancy by their respective TFs may contribute to the activation to different extents. This adds additional freedom to the model, in the form of one extra tunable parameter per TF. The mathematical formalisms of the two mechanisms outlined here, called AB (Activation by Binding) and AP (Activation with Potency), are illustrated in **Figures 3A, B** and explained in Methods.

**Figure 3:**
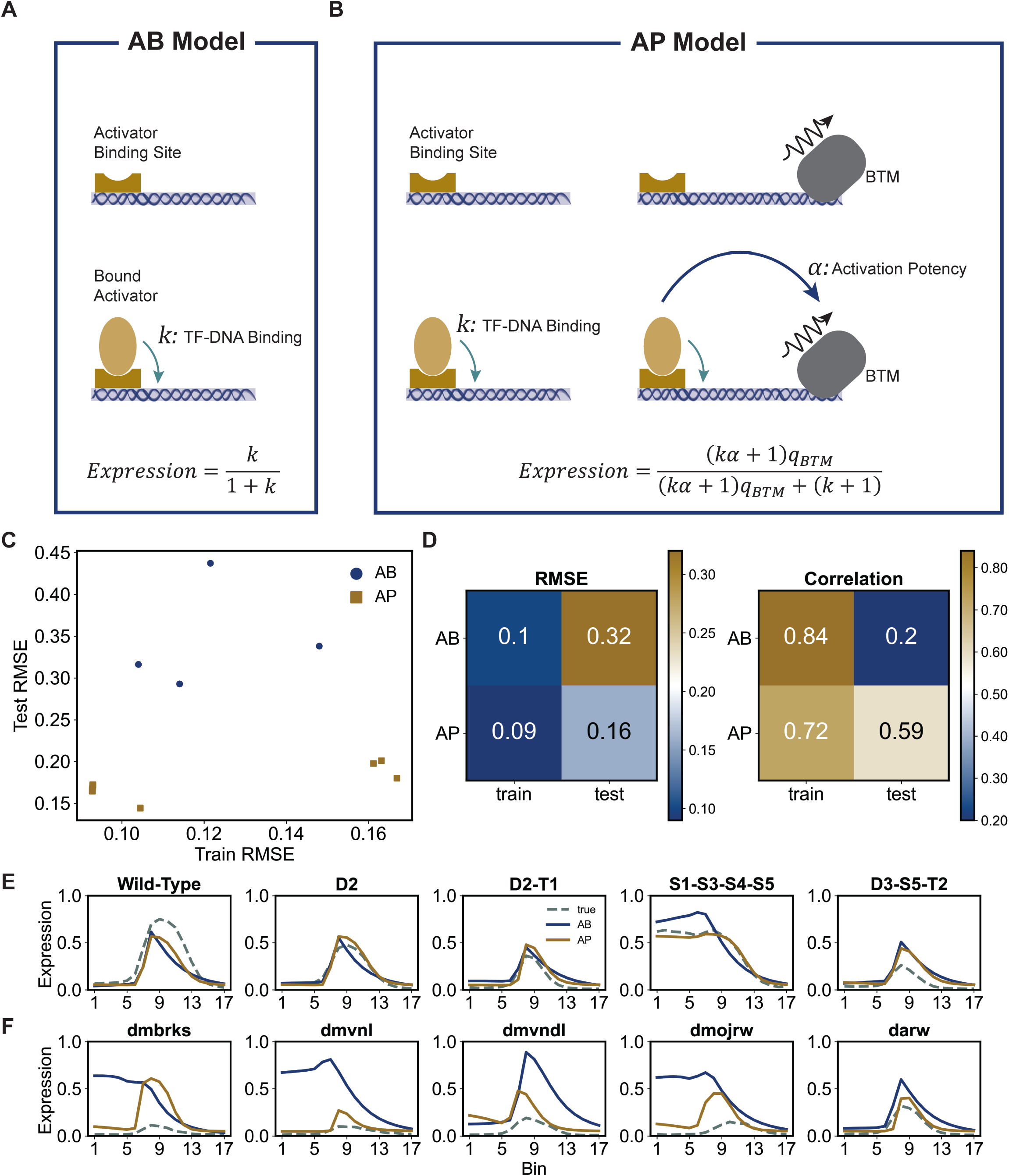
Evaluations of activation mechanisms. (A) Schematic representation of AB model for an enhancer containing only one binding site for an activator TF. Shown are the two possible configurations for this enhancer depending on whether the activator is bound or not. The statistical weights (relative probabilities) of the two configurations are *k* (bound) and 1 (not bound), and expression is proportional to probability of the bound configuration. (B) Schematic representation of AP model for an enhancer containing only one binding site for an activator TF. Shown are the four possible configurations for this enhancer depending on whether the activator and the BTM are bound (or not). Expression is proportional to total probability of the two configurations in which BTM is bound. (C) Train and Test RMSE for the ensemble of 100 AB and 100 AP models. (There is extensive overlap of models (points) in each ensemble.) (D) RMSE and correlation scores of best-fit AB and AP models in train and test data. (E) Predictions of best-fit AB and AP models on select train enhancers (solid curves) are shown along with true expression profiles (dashed curves). (F) Predictions of best-fit AB and AP models on select test enhancers (solid curves) are shown along with true expression profiles (dashed curves).

We sought to determine if the AP and AB models differ in their ability to explain the data set. In the AP model, an activator’s potency was modelled by stipulating an interaction between a DNA-bound activator and the basal transcriptional machinery (BTM), as in GEMSTAT. (This interaction is represented by a single free parameter per TF.) Other mechanistic details such as cooperative binding, short-range repression, etc. are identical between the two models. For a rigorous comparison, we trained ensembles of 100 AB models and 100 AP models on the train set of 38 enhancers and evaluated them on the test set of 13 enhancers. **Figure 3C** shows the accuracy of all models on train and test sets and **Figure 3D** reports on the best-fit model (smallest train error) in both ensembles. In Figure 3D we note that the AP model achieves lower error on the training set, which is not surprising given that it has additional parameters (TF’s potency and basal transcription level). More importantly, it achieves a substantially lower error (RMSE of 0.16 versus 0.32) and higher correlation (correlation of 0.59 versus 0.20) on the test set than the AB model, where the additional parameters do not confer an advantage. This performance gap on test data is apparent at the ensemble level also (Figure 3C), suggesting that it is not an artifact of the optimization step. For both models, the test error values are overall higher than training error, but this is likely to be because the test enhancers are biologically more distinct from the training enhancers than the variation within the training set. The gap between training and test errors is much smaller for the AP model, which has more free parameters, which is contrary to what we would expect if the gap was primarily due to overfitting.

A few illustrative examples of our evaluations are shown in **Figure 3E, F**, where each panel compares AP and AB model predictions to the real expression profiles. While both models exhibit similar accuracy on training enhancers (Figure 3E), the AB model predicts ectopic expression in the ventral region (bins 0-5) for test enhancers ‘dmbrks’ (*D. melanogaster* enhancer of gene *brk*) and ‘dmvnl’ (*D. melanogaster* enhancer of gene *vn*), while the AP model correctly predicts the neuroectodermal peak (bins 7-10) of expression for all shown test enhancers (Figure 3F, Supplementary Figure S2). In light of the above observations, we infer that the data set supports the AP model over the AB model, arguing for separate “potency” for each activator TF, beyond its DNA-binding strength, as an important aspect of the underlying activation mechanism.

#### Repression mechanisms

Previous studies have shown that SNA is a “short-range” repressor whose effect is mediated by a co-repressor named CtBP (43, 44). CtBP can bind to histone deacetylases, which in turn causes DNA to wrap around the histone more tightly and prevents nearby TFs from binding to DNA (45, 46). It implies that a DNA-bound repressor co-bound with CtBP is likely to be the only recruited TF within a small window (∼100 bp) around its binding site. Such a mechanism for short-range repression is implemented in GEMSTAT (28), and we will refer to it as “neighborhood remodeling” (“NR”, **Figure 4B**) below. In this model, the bound repressor makes the neighboring chromatin (within 100 bp in our tests) inaccessible for activators to bind at. An alternative formulation of short-range repression is the “quenching” mechanism (“Q”, **Figure 4A**) (14, 18), which states that a bound repressor will diminish or “quench” the effectiveness of an activator bound nearby; in the thermodynamic model this is achieved by a decreased equilibrium probability of the configuration where both the activator and repressor are bound (see Methods). We tested if the data set can discriminate between the NR and Q models of short-range repression using their implementations within the GEMSTAT framework. We also tested a model with so-called “direct” repression (“DIR”, **Figure 4C**), which, unlike NR and Q, is not a short-range mechanism. Here, the regulatory effect of a bound repressor is due to interactions with the BTM (similar to how activation is modeled in the AP model), and the binding site does not have to be within a short range of any activator binding site for its repressive effect to materialize. Although literature evidence points to short-range repression by SNA (18), we considered it worthwhile to test the direct repression model as a simple phenomenological baseline against which more realistic short-range repression models may be compared in light of the available data. DIR and NR are the least complex of the three models, modelling repression using three free parameters for SNA (DNA-binding, repression potency, and homotypic cooperativity), while Q uses four free parameters (see Methods).

**Figure 4:**
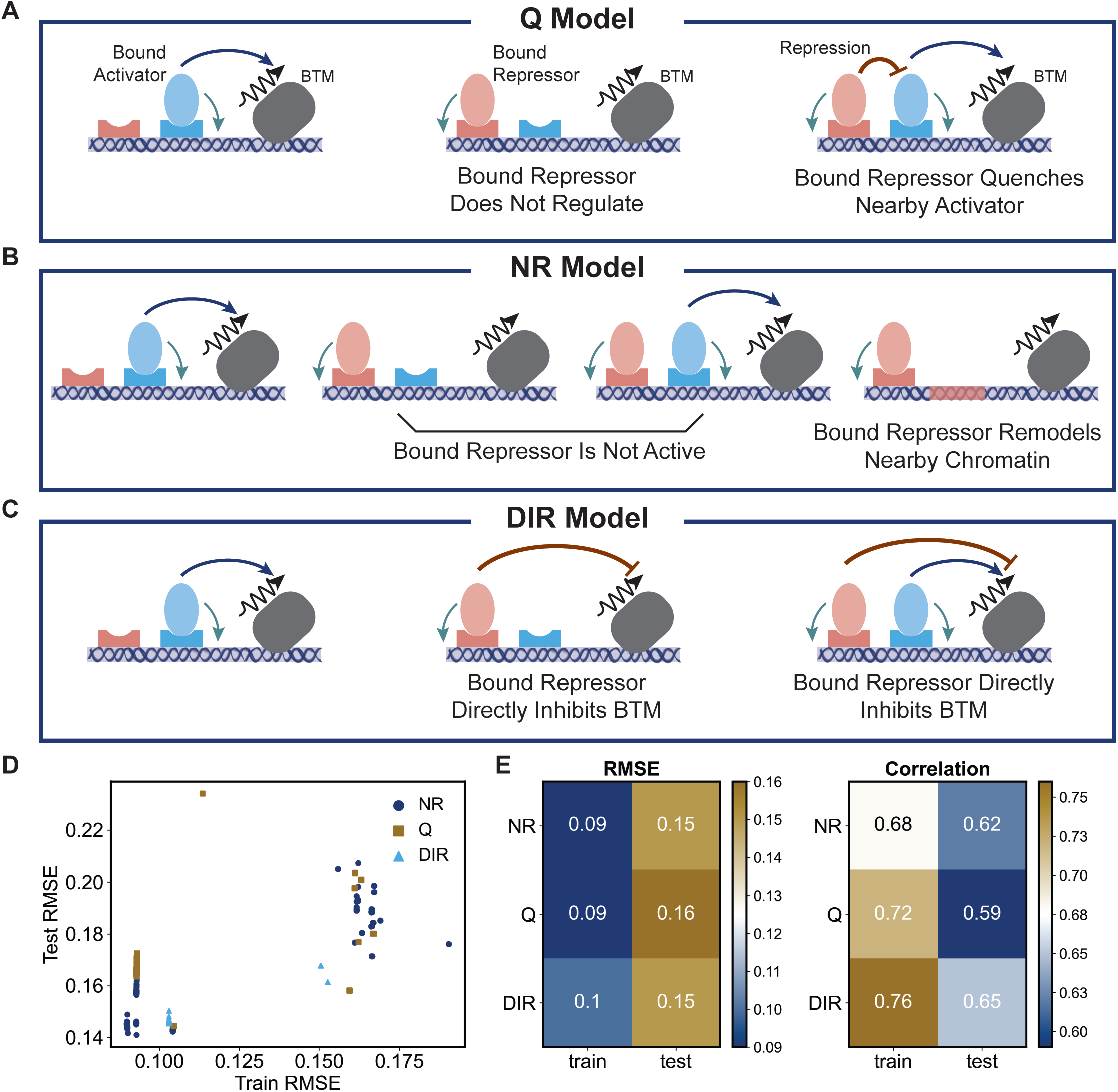
Evaluations of repression mechanisms. (A) Schematic representation of Q model that employs short-range quenching repression mechanism. A bound repressor diminishes the activity of nearby bound activators. (B) Schematic representation of NR model that employs short-range neighborhood remodeling repression mechanism. A bound repressor can be in either active or inactive state. An inactive bound repressor does not interfere with the binding of activators to nearby regions or with the activity of the nearby bound activators, while an active bound repressor prevents activators from binding to nearby regions. Any configuration with an active bound repressor and an activator bound nearby is considered invalid. (C) Schematic representation of DIR model that employs direct repression mechanism. A bound activator directly diminishes the activity of recruited BTM but does not interfere with the binding of activators. (D) Train and Test RMSE for the ensemble of 6000 models for each of Q, NR, and DIR models. (E) RMSE and correlation scores of best-fit NR, Q, and DIR models in train and test data.

**Figure 4D** compares the performance of trained ensembles of NR, Q and DIR models and **Figure 4E** summarizes the performance of the best-trained model in each ensemble. We found both short-range repression models (NR and Q) to yield comparable fits and predictive ability, with the NR model being slightly better in terms of both correlation (0.62 vs 0.59) and RMSE (0.16 vs 0.15) on test enhancers (with one less free parameter than NR model). Interestingly, predictions of the DIR model are as accurate as NR – they show the same RMSE and better correlation on test enhancers, and a significantly better correlation (0.76 vs. 0.68) on train enhancers while using the same number of free parameters. Moreover, correlation values on test enhancers are substantially better with the DIR model than with the Q model. This suggests that in addition to short-range repression, SNA may have long-range repressive effect on transcription of *rho*. We revisit this hypothesis below when we discuss the predictions of our Neural Network-based model.

#### Cooperative activation mechanisms

An important mechanism studied in the context of enhancer function is that of cooperative DNA binding by multiple TFs at proximally located binding sites (2), which results in a synergistic effect greater than the sum of the individual site contributions (11, 42, 47). To test if such effects are reflected in the data, we compared a version of GEMSTAT that models cooperativity between activators (“COOP” model, **Figure 5B**) with one that does not (“NO-COOP”, **Figure 5A**). To implement cooperativity, GEMSTAT includes TF-TF interaction energy terms in configurations where two TFs are bound within a certain distance (set to 50 bp here) of each other. Such terms were presumed for DL-DL, TWI-TWI, and DL-TWI interactions, each represented by a free parameter, in the COOP model. (Both tested versions use the NR model for repression and include homotypic interaction for the repressor SNA.)

**Figure 5:**
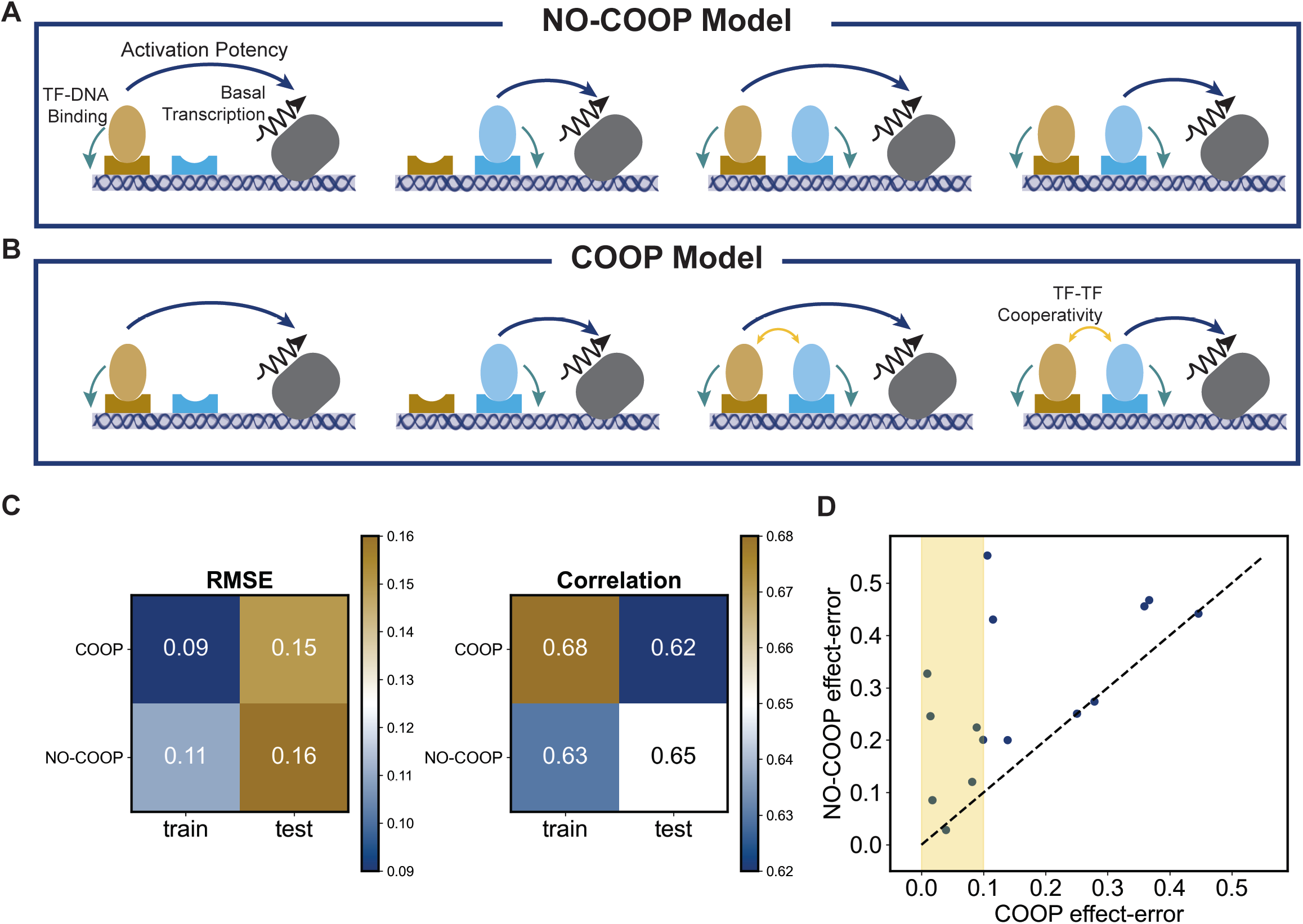
Evaluations of synergistic activation through cooperative binding. (A, B) Schematic representation of the difference between COOP (B) and NO-COOP (A) models. Shown are four of the possible configurations where at least one activator is bound and the BTM is bound. In the COOP model, the two configurations where both activators are bound have favorable interaction between them, resulting in higher probability of these configurations and hence increased expression. (Both COOP and NO-COOP employ short-range neighborhood remodeling repression mechanism; not shown.) (C) RMSE and correlation scores of best-fit COOP and NO-COOP models on training and test data. (G) Effect-error of best-fit COOP versus NO-COOP models on the perturbed enhancers containing deletions of activators’ binding sites (similar to the enhancers shown in Figure 1D). Effect error is defined as the absolute difference between the predicted and true effect of the perturbation; effect of a sequence perturbation (activator site deletion) is defined as decrease in peak (bin 8) expression due to the perturbation. Highlighted region shows perturbed enhancers for which COOP has a small effect error (close to the average training RMSE).

We trained ensembles of models for COOP and NO-COOP separately. The best-trained COOP and NO-COOP models show train error of 0.09 and 0.11 and test error of 0.15 and 0.16 respectively (**Figure 5C**), thus providing some evidence in favor of the former. To tease apart their differences further, we used a complementary evaluation metric: the ability of trained models to predict the effects of activator site deletions (Figure 1D). We defined a model’s “effect-error” as the difference between the predicted effect of the site deletion(s) represented by an enhancer and the real effect captured in the data set. (Here, “effect” is calculated in the same way as in Figure 1D, i.e., the difference in peak expression between wild type and variant enhancer.) **Figure 5D** compares the effect-error of the best-trained COOP and NO-COOP models on each enhancer shown in Figure 1D, where one or two activator sites in the wild type *rho* enhancer have been mutagenized. For all enhancers, the effect-error in COOP is smaller or as low as that in NO-COOP. In particular, in the highlighted region where the effect-error of the COOP model is relatively small (smaller than 0.1), we note several enhancers where the NO-COOP model’s effect-error is substantially worse. These results support the hypothesis that cooperative DNA-binding at proximally located activator binding sites plays a significant role in regulatory function of the *rho* enhancer.

### CoNSEPT: a neural network model of enhancer function

For our final modeling of the data set, we implemented a model called CoNSEPT (**Co**nvolutional **N**eural Network-based **S**equence-to-**E**xpression **P**rediction **T**ool), that can accommodate highly non-linear contributions of TF binding sites to overall enhancer function (**Figure 6**). Our primary goal was to test if a convolutional neural network (CNN) model, which does not explicitly incorporate known rules of cis-regulatory encoding, can learn such rules (regulatory mechanisms, including distance-dependent interactions between sites) from the data. This tool first uses pre-determined TF motifs (PWMs) to scan both strands of an enhancer to identify putative binding sites and estimate their strengths (“PWM scores”). It then integrates these strengths with respective TF concentration values to assess each TF’s presence at varying locations in the enhancer, analogous but not identical to occupancy in thermodynamics-based models (see Methods). Next, it assembles the presence scores of each TF pair into a feature matrix, which is passed to a separate convolutional filter to capture short-range interactions between the TF pair. Outputs of these filters are aggregated and passed to the subsequent convolutional layers to capture long-range interactions. Finally, the output of the last convolutional layer is combined linearly to predict the expression driven by the enhancer (see Methods for details).

**Figure 6:**
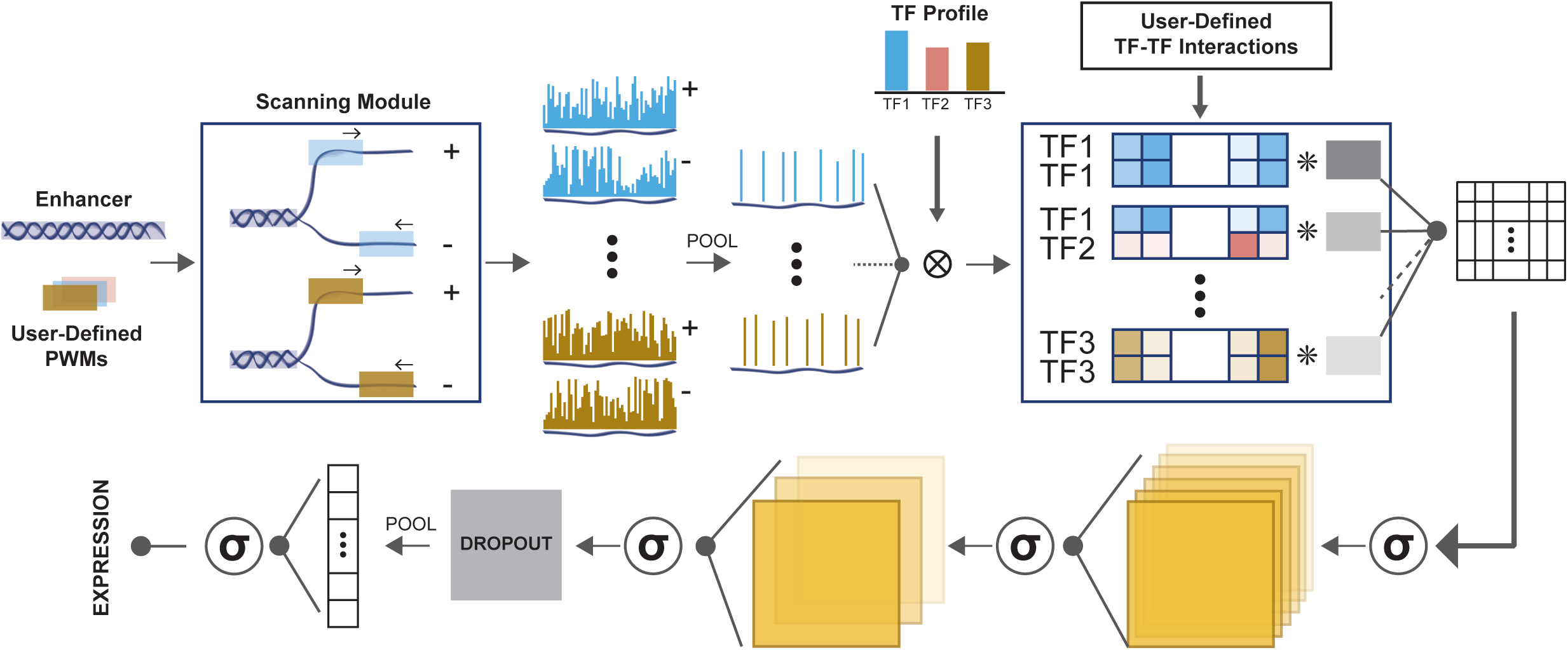
Architecture of CoNSEPT. First the input enhancers are scanned with user-defined PWMs and passed to a pooling layer to extract the strongest matches. Next, TF profiles are integrated (multiplied) with the extracted scores and pair-wise features are built according to the user-defined TF-TF interactions. Each pair-wise feature is passed to a separate kernel capturing short-range regulatory interactions between pair of TFs. (Note: a TF pair may be homotypic.) The concatenated output of interaction convolutional kernels is passed to a sequence of activation functions (*σ*) and optional additional convolutional layers. The final activated output of the convolutional layers is passed to a dropout layer followed by a fully-connected layer and an activation function to predict the expression.

We used the training data comprising the WT *rho* enhancer and its 37 variants to train CoNSEPT models; eleven of the 13 enhancers in the previously defined test data were used for evaluating trained CoNSEPT models. (Two enhancers were removed from the original test data due to their significant length difference with the training enhancers; see Methods.) We used data on three additional enhancers (variants of the *rho* enhancer, regulated by the same TFs) available from Sayal et al. (14) as a separate “validation set” for tuning hyper-parameters of CoNSEPT models. Previous applications of neural networks for regulatory sequence interpretation had the benefit of large training data sets (37, 39), and the challenge for us was to train a CNN on a far smaller data set without “overfitting”.

An ensemble of CoNSEPT models with varying architectures and hyperparameters (see Supplementary Table S4 for different settings used) was trained on training data and evaluated on validation data (**Figure 7A**). A subset of the ensemble exhibits better training and validation accuracy than the best-trained GEMSTAT model (NR repression, COOP), and we selected among these the one with the smallest validation error (circled) for further analysis (see Supplementary Table S5 for the settings of this model). **Figure 7B** compares the RMSE and correlation values for the predictions of this model on train, validation and test data sets with predictions of GLM and GEMSTAT models learned above as the best representatives of the linear and thermodynamics-based models. CoNSEPT predictions on the test set are at least as accurate as GLM and GEMSTAT in terms of RMSE, and moderately better in terms of correlation values (Examples of its improved prediction are shown in **Figure 7C, D**.) Its high accuracy values on the training set are not surprising given its higher number of free parameters (1537, 11, and 4 for CoNSEPT, GEMSTAT and GLM models respectively), but its competitive test accuracy suggests that despite the vastly greater model complexity and limited training data, ConSEPT model fitting does not suffer from overfitting any more than GLM and NR models do.

**Figure 7:**
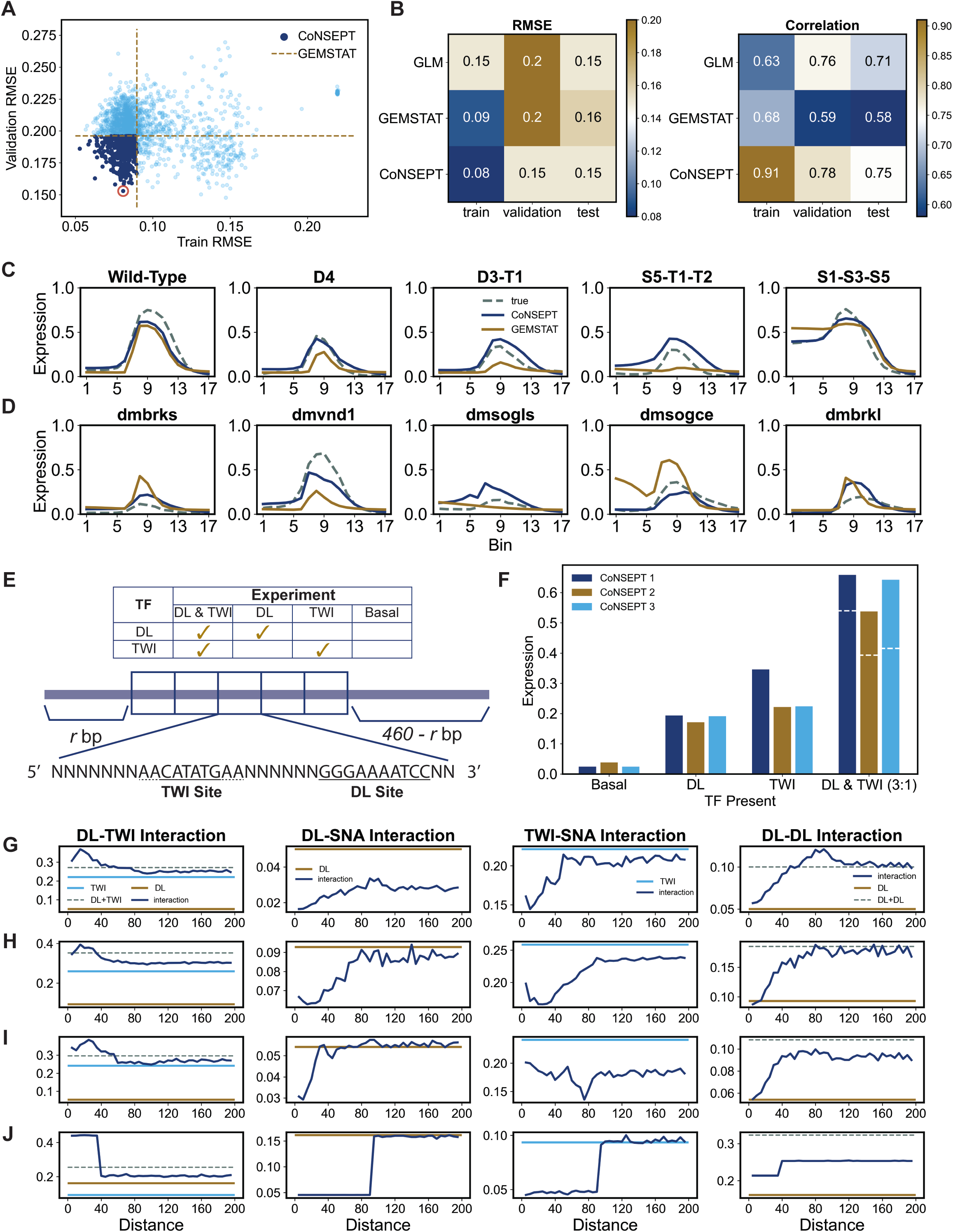
Evaluations of CoNSEPT’s predictions. (A) Train and validation RMSE for the ensemble of CoNSEPT models. Best-fit GEMSTAT (NR, COOP) model was used as a baseline (dashed lines). Among the CoNSEPT models with a better performance than the baseline, we selected the one with the smallest validation RMSE (circled). (B) RMSE and correlation scores of the selected CoNSEPT model on training, validation and test data. (C, D) Predictions of the selected CoNSEPT model and the baseline GEMSTAT model on select train (C) and test(D) enhancers for which CoNSEPT improves predictions. (E) In silico synthetic construct adopted from (42) consists of 5 repeating blocks each containing one DL and one TWI binding site with a length of 35 bp. The 5-block construct with a length of 175 bp is randomly padded by r (random variable) “dummy” bases on the left and 460-r dummy bases at the right end to get a longer construct of length 635 bp consistent with the length of train enhancers. The top table shows the four trans contexts for which this construct’s expression was predicted: “DL & TWI” where both TFs are present with a 3:1 concentration ratio, “DL” where TWI concentration is zeroed out, “TWI” where DL concentration is zeroed out, and “Basal” where neither of the factors are present. (F) Predictions of the three selected CoNSEPT models on the synthetic construct shown in E. The white dashed lines on DL & TWI bars correspond to the sum of the expression driven by DL and TWI individually. (G-J) Predictions of the three selected CoNSEPT models (G-I) and the baseline GEMSTAT model (J) on constructs containing only one or two TF binding sites with different inter-site spacings (0-200 bp; x-axis). Each column corresponds to a pair of interacting TFs. The dark blue curves show predicted expression on constructs containing two binding sites for the TF pair. The brown and light blue curves show predicted expression on construct with one DL site or one TWI site respectively. Dashed lines show the expression obtained by summing the predicted expression driven by individual factors (DL+TWI in first columns and DL+DL in fourth column).

### CoNSEPT learns distance-dependent interactions between TF binding sites

We saw above that CoNSEPT learns to predict expression from sequence with almost no pre-determined notion of cis-regulatory grammar such as activation, repression or pair-wise interactions between sites. We next interrogated the trained model to determine if biophysically plausible cis-regulatory rules underlie its highly parameterized non-linear form. We were particularly interested in whether it learns distance-dependent pairwise interactions between TF binding sites. Recall that the linear models above do not allow such interactions and our thermodynamics-based formulations must be “hard-coded” with a particular form of distance-dependent interaction.

Prior to interrogating the CoNSEPT model, we refined it based on an additional data set that specifically captures synergistic interactions between DL and TWI binding sites. Shirokawa et al. (42) tested a synthetic enhancer consisting of 5 repeating blocks of a sequence with DL and TWI binding sites six bp apart (**Figure 7E**); a reporter assay was used to show that expression driven in the presence of both DL and TWI (in 3:1 ratio) is greater than the sum of expression by these TFs separately, suggesting synergistic activation by these two TFs (42). Our model refinement demanded that the CoNSEPT model with architecture and hyper-parameter setting as determined above (circled model in Figure 7A) additionally recapitulate the synergistic activation encoded by the construct of Shirokawa et al. (We also required expression driven by TWI alone to be at least as high as that driven by DL, another observation made by the authors.) This refinement step yielded three CoNSEPT models (parameterizations), whose predictions on the synthetic enhancer of Shirokawa et al. are shown in **Figure 7F**. (See Methods for details of refinement steps.)

We next used the three models derived above to predict the expression driven by constructs harboring a single pair of TF binding sites (DL-DL, DL-TWI, DL-SNA, or TWI-SNA) at varying inter-site distances (see Methods). The results, shown in **Figure 7G-I**, reveal the distance-dependent interactions between TF binding sites, as learnt by CoNSEPT models. Firstly, all three models predict synergistic activation by a DL-TWI site pair (first column) over short distances < 40 bp with a maximum effect at ∼20 bp inter-site spacing consistent with previous experimental findings (11). The GEMSTAT model (NR, COOP), shown in the bottom row for comparison, shows a similar trend; indeed, GEMSTAT training required that cooperative binding be of a fixed strength within 50 bp and absent for greater inter-site spacing. On the other hand, the trend was learnt by CoNSEPT models entirely from training data. Notably, the DL-TWI spacing has been shown to be a key element of functional organization of neuroectodermal enhancers (48).

We next examined learnt distance-dependencies for DL-SNA interactions (Figure 7G-I, second column). All three models correctly predict SNA as a repressor, with two predicting its effect to be exclusively short-range, decreasing linearly as the distance from DL (activator) site increases from 0 to 40 bp or 80 bp. The third model (CoNSEPT 1, Figure 7G) also shows the 80 bp short-range effect but predicts an additional long-range effect that extends to the maximum spacing interrogated. The GEMSTAT model (bottom row) is consistent with a short-range effect, as observed above (Figure 5), but its distance-dependence function (range of 100 bp) was hard-coded into the model. Interestingly, our GEMSTAT modeling had also indicated the possibility of longer-range effects of SNA (‘DIR’, Figure 5), akin to that seen with the CoNSEPT 1 model. Similar trends were seen with TWI-SNA interactions, with two of the models predicting a short-range effect (range of 60 or 90 bp) (Figure 7G, H, third column) and one model (CoNSEPT 3) also indicating a longer-range effect. The consistent observation of long-range (beyond 100 bp) repressive effects of SNA sites, through CoNSEPT models (DL-SNA as well as TWI-SNA interactions in Figure 7G-I) as well as GEMSTAT models (Figure 5), suggest that mechanisms other than the documented short-range repression by SNA may be at play in neuroectodermal enhancer function.

Intriguingly, all three CoNSEPT models predict (Figure 7G-I, fourth column) that the activation by two DL sites is less than twice the activation by a single site at shorter inter-site spacing (< 50 bp). The GEMSTAT model’s predictions are similar (bottom row) – when asked to tune a parameter representing DL-DL interaction (limited to sites within 50 bp), the model learnt an antagonistic interaction. The models mostly did not find evidence of synergistic interaction between DL site pairs at any spacing. In fact, the CoNSEPT models predict that two adjacently placed DL sites drive no more activation than either site alone, while the GEMSTAT model, forced to assume fixed interaction strength within the distance threshold (50 bp), differs in this prediction. We do not know of a plausible candidate mechanism for the prediction of antagonistic DL-DL site interactions at short distances, and the finding needs more direct confirmation and dissection in the future.

## Discussion

Sequence-to-expression modeling can reveal mechanistic details of an enhancer’s regulatory function as encoded in its sequence. The significance of such modeling is well argued in the literature (16) and, with rapidly advancing technology for multiplexed assays of enhancer function (49–51), a rigorous method for mechanistic inference and cis-regulatory decoding is clearly needed. We were interested in a version of this problem where the model captures variation of function across different enhancers as well as across cellular conditions; the latter requires that the model utilize cellular TF concentrations in making predictions. Fortunately, prior work offers several ways forward, from linear models (35, 40) to thermodynamics-based models (14, 18, 21, 28, 52, 53). Each of these modeling approaches has been shown to explain one or more expression data sets and reveal useful insights into their underlying biology. Typically, each approach has its own mechanistic assumptions baked into the implementation, with quantitative details of assumed mechanisms being left as trainable parameters. To our knowledge, no study has attempted to fit models of fundamentally different form, i.e., with different mechanistic assumptions (or lack thereof), to the same data set to assess their relative potential to explain the data and provide mechanistic clues. This existing gap was the primary motivation behind our work.

We tested linear and generalized linear models, thermodynamics-based models with varying biophysical assumptions, as well as a newly implemented convolutional neural network on the same data set, evaluating their ability to fit the data and generalize to unseen (but related) enhancers. We found these different modeling approaches to show similar predictive ability overall in terms of the RMSE score, although the neural network model (CoNSEPT) yields higher correlation coefficient on average (Figure 7B) compared to other models (See supplementary table S6). We found that linear models, which are simple to implement and use, provide good fits to data but are by construction unable to discover non-linear combinatorial contributions of multiple regulators. On the other hand, equilibrium thermodynamics-based models allow incorporating various hypotheses regarding combined action of multiple regulators in ways that depend on inter-site distance.

In evaluating different thermodynamics-based frameworks (14, 28), we found that modeling the effects of activators via two separate interactions (TF-DNA binding and TF-BTM interactions) provides a substantial benefit in terms of predictive power (Figure 3), compared to modeling activator-DNA binding alone, at least within the context of our evaluations. The choice of separating the DNA-binding aspect of a TF’s function from its expression activation aspect is not arbitrary – these two aspects are typically encoded in different domains of the protein, and such separation is common when modeling repressor TFs (18, 25). Activator potency is often attributed to interactions between TF and the basal transcriptional machinery (potentially mediated by co-factors), whose details are expected to differ from one TF to another. Note that we used the implementation of Sayal et al. for the AB model but set its TF-TF interaction rules to a simpler form than those of the original study (14), in order to match those in the AP model (implemented in GEMSTAT). We expect that replicating the more complex settings of interaction rules of Sayal et al. will yield better fits to the data, but make direct comparison (of AP and AB) challenging.

We tested several additional thermodynamics-based formalisms within a common framework (GEMSTAT), using the same parameter training algorithms. We saw evidence of cooperative DNA-binding at proximal sites by TF pairs (Figure 5), as has been reported extensively in the literature (2), including through thermodynamics-based models (28, 54). Importantly, DL-TWI synergistic action has been experimentally observed (11, 42). When testing three different modes of repression, we were surprised to note that a “direct repression” mechanism, where a bound repressor’s regulatory contribution does not depend on how far its binding site is from an activator’s site, has the same predictive ability as the two “short-range repression” mechanisms tested. In light of literature evidence for short-range repression (55, 56), this finding suggests that discerning true mechanisms from modeling of experimental results *ex post facto* may not be adequately powered, and that active learning approaches (57, 58) that suggest the most informative future experiments may be called for.

While thermodynamics-based models allow interactions between TF binding sites to depend on their relative arrangement in the sequence, the form of the dependence has to be pre-determined. For example, commonly, cooperative DNA binding and short-range repression are assumed to have a fixed strength as long as inter-site distance is less than a pre-set threshold (18, 21, 25) and absent otherwise. This assumption reduces the complexity of the model, though it may oversimplify the distance-dependence of TF interactions. (Sayal et al. (14) considered a more flexible form for this dependence, with different “bins” of inter-site distance being associated with tunable interaction strengths.) Moreover, typical thermodynamics-based models allow interactions only in configurations where the two TF bound to proximal sites have no other bound TF in between; this is necessitated by considerations of computational efficiency. These constraints of thermodynamics-based models led us to encode a more flexible model of distance-dependent TF-TF interactions (cooperative as well as antagonistic) in CoNSEPT, through layers of convolution kernels that process all putative site pairs near each other. By examining the trained model’s predictions on a simple synthetic enhancer with varying inter-site distances, we were then able to extract the distance-dependence function learned by it (Figure 7G-J) in a data-driven manner. The learned function is mostly in agreement with that used in the thermodynamics-based models, e.g., both approaches suggest cooperative effects of DL-TWI site pairs within 40 bp of each other (11), as well as repressive effects of SNA sites located within 80 bp of an activator site. Interestingly, one of the three CoNSEPT models is consistent with “direct repression” (not limited to short-range effects), an observation made also when fitting thermodynamics-based models with this form of repression.

A key contribution of this work is the design and implementation of CoNSEPT, which is a general-purpose tool for sequence-to-expression modeling using convolutional neural networks. Our work demonstrates the feasibility of training such complex models (thousands of free parameters) on limited data sets (hundreds rather than thousands of samples), and we have tested that it can handle other data sets of much larger size (tens of thousands of samples, data not shown). It stands in contrast to other available implementations of neural networks for regulatory genomics, which are targeted to modeling epigenomic (39, 59, 60) and cistromic (36, 38) data, or do not explicitly model the dependence of sequence function on cellular descriptors such as TF levels (61). This feature allows CoNSEPT to make predictions for varying cellular conditions. Moreover, its reliance on pre-defined PWMs is different from previous applications of neural networks in regulatory genomics where TF motifs were learned from the data using convolutional kernels (38, 39, 60). We believe that using known PWMs, which are often reliably characterized experimentally (62), reduces the number of free parameters and lowers the chance of overfitting.

The work most closely related to the CoNSEPT model is the neural network model presented by Liu et al. (63) who used it to model the expression driven by enhancers related to the *eve* gene of Drosophila. Their model is a recasting of the thermodynamics-based model of Kim et al. (54) and includes all of the latter’s mechanistic aspects. At the same time, the model of Liu et al. is specialized for the biological system (*eve* enhancers) studied by them and encodes distance-dependence of interactions with pre-determined form and parameters motivated by that system. CoNSEPT, on the other hand, departs from the thermodynamics-based formalism and takes a more data-driven approach to capturing cis-regulatory “grammar” and distance-dependent interactions.

In summary, our work shows that there are multiple formalisms capable of explaining the sequence-function mapping encoded in enhancers, with complementary strengths and varying reliance on prior mechanistic knowledge. Although the mechanisms that we find are specific to the regulation of *rhomboid* and other neuroectodermal enhancers, and might not necessarily generalize to the regulation of other genes, our work gives a recipe for understanding the regulatory mechanisms in a data-driven or model-driven manner. In addition to their explanatory role, the models tested here can also be useful for predicting the expression driven by unseen sequences and cellular contexts; this ability has several applications in down-stream analysis such as predicting the effect of a particular TF’s knockout, site mutagenesis, or effects of single nucleotide polymorphisms on the expression (64, 65). We also showed that convolutional neural networks can be a reliable expression prediction tool capable of learning non-linear regulatory mechanisms from modest-sized training data.

## Methods

### Linear models

#### LM model

Expression (*E*) driven by an enhancer is a weighted sum of contributions from TFs that bind to the enhancer:

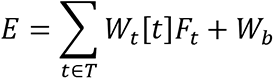

where *T* is the set of TFs, *W*_*t*_ is a TF-specific weight that reflects its activating or repressive role and strength, *W*_*b*_ is a “basal” expression parameter, [*t*] is the concentration of TF *t,* and *F*_*t*_ reflects the total binding site presence of *t* on the enhancer, defined as:

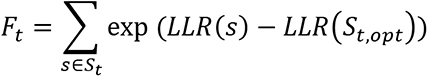

where *S*_*t*_ is the set of all putative binding sites of *t* in the enhancer, *S*_*t*, *opt*_ represents the strongest possible binding site of *t* and *LLR*(*x*) denotes the log likelihood score of site *x*, calculated using the given Position Weight Matrix (PWM) of *t* and a provided background nucleotide distribution. This definition of site strength follows (66).

#### GLM model

Expression driven by an enhancer is a sigmoid function of the LM model:

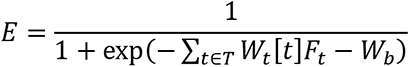

#### GLMQ model

This defines the total contribution of TFs as a non-linear function of their concentration [t] multiplied by their site strength *F*_*t*_ and then applies a sigmoid function to model saturation:

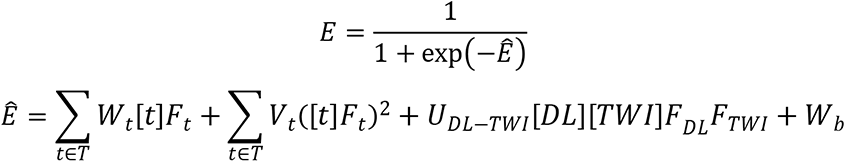

where *V*_*t*_ is a free parameter associated with *t* − *t* cooperative interaction. In addition to linear terms [*t*]*F*_*t*_ and quadratic terms ([*t*]*F*_*t*_)^2^ for each TF, this includes a term for heterotypic cooperativity of DL and TWI, as suggested in the literature (14, 42), with a tunable weight *U*_*dl*−*twi*_.

### GEMSTAT model

All of the thermodynamics-based models explored in this work are implemented in GEMSTAT (28), except the AB model, for which the implementation of Sayal et al. was used (14). Thermodynamics-based modeling of gene expression involves enumerating all “microstates” (henceforth, states) of the enhancer under thermodynamic equilibrium. A state is defined as a configuration specifying the bound or non-bound status of each TFBS. Therefore, an enhancer containing *n* TFBSs has 2^*n*^ states. In GEMSTAT, for each of these states the Basal Transcriptional Machinery (BTM) may be in the bound or non-bound state, making 2^*n*+1^ states. We define “ON” states of the system as those where BTM is bound; other states are called “OFF” states. The expression driven by the enhancer is assumed proportional to the probability of the system being in ON state:

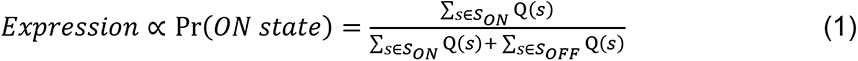

where *S*_*ON*_ and *S*_*OFF*_ denote the set of all ON and OFF states of the enhancer, respectively, and Q(*s*) is the Boltzmann weight that prescribes the relative probability of state *s* in equilibrium. (The denominator is the partition function.) For simplicity, we set the constant of proportionality in the above equation to 1.

The Boltzmann weight of state *s*, Q(*s*), is calculated as the product of terms representing each molecular interaction in that state; these interactions include TF-DNA interactions at binding sites, BTM-DNA interaction at promoter, TF-BTM interactions representing activation or repression effects of TFs, and TF-TF interactions representing cooperativity or antagonism between proximally DNA-bound TF pairs. See supplementary table S3 for the parameters used in different GEMSTAT models in this study.

#### TF-DNA interaction

The weight (term contributed to Q(s)) of a TF-DNA interaction at site *s* for a TF *t* is given by:

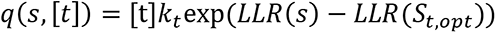

where *k*_*t*_ is a TF-specific free parameter that is learned from the data and other terms are as defined above.

#### TF-BTM interaction

The weight (term contributed to Q(s)) of a TF-BTM interaction is a TF-specific positive constant which is > 1 for activators (making the ON state more favorable than corresponding OFF state) and < 1 for repressors. This TF-specific constant is referred to as the TF’s “potency” in the Results section. In this study, repressor’s potency is present only in DIR model representing long-range repression. We employed the GEMSTAT’s “limited contact” scheme of activation, where at most one bound activator can interact with the BTM in any state.

#### TF-TF interactions

Any activator-activator or repressor-repressor pair (i.e., DL-DL, TWI-TWI, SNA-SNA, DL-TWI) bound within 50 bp of each other is modeled as interacting, with the weight contributed to Q(s) being a learnable free parameter. Such interactions may be configured to be excluded from or included in the model e.g., when comparing the “COOP” and “NO-COOP” settings of GEMSTAT. Separately, an activator-repressor pair (i.e., DL-SNA, TWI-SNA) bound within 100 bp of each other is modeled as interacting, with a learnable weight (<= 1). Such interactions represent short-range repression by quenching, following Sayal et al (14), and may be configured to be excluded or included in the model.

*BTM-DNA interaction* is modeled as a single learnable parameter.

### Activation by Binding (AB) model

We used the implementation of Sayal et al. (14) for the AB model. Here, the bound or non-bound status of the BTM is not part of the state definition, and expression is assumed proportional to the probability of states with at least one bound activator that is not repressed by a bound TF nearby. The Boltzmann weight of a state is defined as the product of terms representing TF-DNA and TF-TF cooperative interactions, defined as in GEMSTAT. Terms involving BTM interactions are not part of the model and in particular TF-BTM interaction parameters that capture activating and repressive influences in the GEMSTAT model are not included. See (14) for details. We used the “binned” interaction scheme of their model, using one bin and setting the ranges for TF-TF interactions to match those of GEMSTAT (50 bp for cooperative interactions, 100 bp for short-range repression).

### Alternative repression mechanisms in GEMSTAT

In the GEMSTAT model used for testing the AP mechanism of activation, repression is modeled by short-range (<= 100 bp) activator-repressor interactions (DL-SNA, TWI-SNA). The repression mechanism is referred to as “Q” (for “quenching”). An alternative to this is the “Neighborhood Remodeling” (NR) mechanism of short-range repression, where any repressor site may be in one of three possible states (rather than two): “non-bound”, “bound-only” and “bound-effective” (28). In the bound-effective state, the bound repressor modifies its neighborhood on the DNA such that the neighboring chromatin becomes inaccessible for other TFs to bind. This modification is assumed to occur within a fixed distance *d*_*r*_ from the bound repressor site. States where a repressor site is in the “bound-effective” state and another site (for any TF) within *d*_*r*_ distance is in the bound state are considered invalid. The “bound-only” state is akin to the usual “bound” state of a site, with no restrictions on possible states of neighboring sites. A site in the bound-effective state contributes an additional factor *β*_*r*_ (a TF-specific free parameter) to the Boltzmann weight of the overall state; a higher value of *β*_*r*_ (> 1) leads to lower fractional occupancy of proximal activator sites, thus achieving greater repression. In this study we used a range parameter of *d*_*r*_ = 100 *bp*. A third alternative to the “Q” and “NR” mechanism is the “DIR” (for “direct” repression) mechanism, modeled by a repressor-BTM interaction regardless of distance of repressor binding site from a bound activator site, thus making this a “long-range” repression mechanism (28).

### CoNSEPT model

CoNSEPT (Convolutional Neural Network-based Sequence-to-Expression Prediction Tool) is a neural network model that predicts the enhancer activity as a function of enhancer sequence and TF concentration levels. The model is parameterized by user-provided PWMs (motifs) representing TF binding preferences.

### CoNSEPT architecture

First, the enhancer sequence (of length *L*) is scanned with user-defined PWMs to score the presence of each motif along the enhancer. The scanning module computes the complementary sequence (negative strand) of the input enhancer and converts both strands into a one-hot encoded representation by replacing each nucleotide (A, C, G, or T) with a 4-dimensional vector as follows:

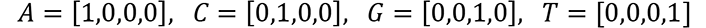

User-defined PWMs are formatted into a set of *K* x 4 matrices, *m*_*t*_ for each TF *t*, representing the probability of each nucleotide appearing at each position of the TF’s binding site of length *K*. In this study, we used *K* = 10, and to do so we had to expand the TWI and SNA PWMs we obtained from Sayal et al. (14) by three and two bases, respectively, with a probability of 0.25 over A, C, G, and T.

Both encoded strands are then scanned with each motif using a convolution operation:

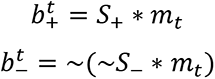

where 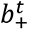 and 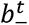 represent the binding score profiles of TF *t* on the positive and negative strands, respectively, *S*_+_ and *S*_−_ denote the positive and negative encoded strands. Moreover, ∼(.) represents a “flip” operation that reverses the sequence. The flip operation mimics how the negative strand is scanned for motif presence in previous work (14, 25, 26, 28), and as far as we know, this is the first time it has been included in a neural network model of sequence-function.

Next, for each TF *t*, the positive and negative binding score profiles, represented by an *L̇* x 2 matrix (*L̇* = *L* − *K* + 1), are passed to a 2-dimensional max-pool layer that extracts the strongest binding site score from the two strands within a window of a certain width. Since we have three TFs (DL, TWI, and SNA) in this study, this step gives us three *L̈* -dimensional vectors *B*^*t*^, where *L̈* < *L̇* is the reduced length due to max-pooling. These vectors are then integrated with the TFs’ concentration values to obtain “occupancy” vectors *F*_*t*_:

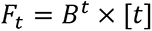

where *F*_*t*_ is the vector representing occupancy of TF *t* along the enhancer, [*t*] denotes the concentration of TF *t* in a particular cellular context, and ‘x’ represents element-wise product.

Next, CoNSEPT incorporates user-specified prior knowledge of TF-TF interactions. To this end, for a specified interacting pair, (*t*_1_, *t*_2_), an *L̈* x 2 “TF pair feature matrix” is constructed by stacking *F*_*t*1_ and *F*_*t*2_. The specified interactions may correspond to homotypic or heterotypic cooperativity or short-range repression. Each TF pair feature matrix is passed to a separate 2-dimensional convolutional kernel that moves along the enhancer length and captures the short-range patterns in occupancies, producing an *L̇̈*-dimensional vector Ω_*i*_ (*L̇̈* < *L̈* is the reduced length due to convolution):

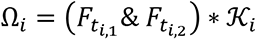

where *i* corresponds to a specified TF-TF interaction between *t*_*i*, 1_ and *t*_*i*, 2_, 𝒦_*i*_ denotes the convolution kernel for this interaction and “&” is the stacking operation. The outputs Ω_*i*_ of all convolutional kernels are stacked into a *L̇̈* x *N* matrix Ω, where *N* denotes the total number of user-defined TF-TF interactions.

In this study, we used DL-DL, TWI-TWI, and SNA-SNA interactions, consistent with the homotypic cooperativities in COOP model, also DL-TWI interaction, consistent with the heterotypic cooperativity in COOP model, and SNA-DL and SNA-TWI interactions, consistent with the short-range repressions in COOP model. The output of the convolutional kernels, Ω, is activated by a non-linear function. We also tested passing this activated output into two additional convolutional layers with different number of kernels and activated by a non-linear function after each layer (Figure 6), in order to capture longer-range interactions. The activated output of the last convolutional layer goes into a dropout layer that is widely used for regularizing neural network models (67). We did not use dropout in the previous layers to maintain any positional feature on the enhancer that might contribute to short-range and long-range regulations. The output of the dropout layer is passed to a non-overlapping max-pool layer that extracts the strongest signals. Finally, the output of this pooling layer is linearly combined through a fully connected layer and goes into a final activation function that outputs the expression value. For this last activation function, we tested sigmoid and tanh functions suitably modified to ensure positive expression values upper-bounded by one.

We use the following naming conventions to fully demonstrate CoNSEPT’s architecture:

***BS*:** TF binding-site scanning module
***PFC*_*α*, *β*_:** Stacking of occupancy vector of each pair of TFs and passing the resulting TF pair feature matrix to a 2-dimensional convolutional kernel of size (*α*, 2) and stride (*β*, 2). This unit generates the output Ω described above. In this study we used *α* = *β*; i.e. non-overlapping convolution.
***P*_*γ*, *δ*_:** a max-pool layer of size *γ* and stride *δ*
***CN*_*k*_:** a 1-dimensional convolutional layer with *k* kernels of size 3 and stride 1.
***FC*:** a fully connected layer
***DR*_*p*_:** a dropout layer with probability of *p*
***LN*:** a layer-normalization layer (68)
***σ*:** an activation function
**× [*TF*]:** multiplication by TF concentration values to obtain occupancy as described above
**Architecture:** *BS* − *LN* − *P*_*γ*, *δ*_ − × [*TF*] − *PFC*_*α*, *α*_ − *σ*_1_ − {*CN*_*k*1_ − *σ*_1_ − *LN*}_*c*_ − *DR*_*p*_ − *P*_3, 3_ − *FC* − *σ*_2_

Above, {.} represents an optional block of convolutional kernels followed by activation and normalization and *c* denotes the number of blocks used in the model. Note that for *c* = 0, the additional convolutional layers inside the braces are not used. In this study, we tested 2016 different models for various settings of *γ*, *δ*, *α*, *σ*_1_, *c*, *k*_1_, *k*_2_, *p*, and *σ*_2_ and selected the best model based on the performance on a validation data set. See supplementary table S4 for the parameter settings tested.

#### Training CoNSEPT

To train a CoNSEPT model we find the settings of parameters *θ* that minimize the mean squared error between the model output (*Ê*) and the ground-truth expression (E) over the training data:

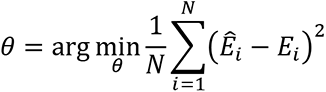

where *N* is the total number of training samples, each representing a different combination of enhancer and cellular context (bin along DV axis). For optimization, we employed a stochastic gradient descent algorithm using Adam optimizer (69) for 1000 epochs and a batch size of 20. The learning rate of the gradient descent was scheduled to be decreased during the training.

#### Synthetic constructs for evaluation of DL-TWI synergistic activation

In order to guide the training of CoNSEPT models towards capturing the cooperative activation by DL and TWI, we used data from Shirokawa et al. (42). We constructed a DNA sequence mimicking that tested by Shirokawa et al., consisting of a TWI and a DL binding site located 6bp apart on a construct of length 35 bp with the following sequence:

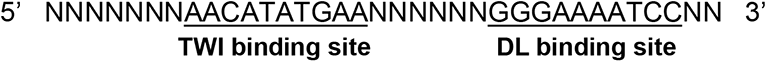

where “N” denotes a dummy base. The DL binding site is slightly different than that used by Shirokawa et al. (42) since we used the consensus DL binding site implied by the PWMs we obtained from Sayal et al. study (14). Also, the TWI site is four bp longer than that in Shirokawa et al. (42) due to the padding of TWI PWM in our study.

Similar to the experiments of Shirokawa et al. (42), we next created a sequence with five consecutive repeats of the above block. Since CoNSEPT was trained on enhancers of length 635 bp (see See supplementary Note S1), we expanded this construct of length 175 bp to 635 bp by adding dummy bases (“N”) at both ends. To eliminate potential biases, we repeated this expansion with 80 different random distributions of dummy bases at the two ends; therefore, we obtained 80 different constructs of length 635 bp that only differ in their distribution of dummy bases at the two ends.

Next, we selected the hyper-parameter settings of CoNSEPT with the highest validation set performance (referred to as “best-validated” CoNSEPT model) among the ensemble of 2016 settings described above. Using these hyper-parameter settings, we re-trained an ensemble of 200 CoNSEPT models on the training data with different random initializations of free parameters. We examined the prediction of each of these 200 trained models on the synthetic constructs defined above, assuming DL and TWI concentrations in a 3:1 expression ratio (similar to that in experiments of Shirokawa et al. (42)) as well as in the presence of only one of the TFs and a basal level in the absence of both TFs. For each model, the predicted expression values were averaged over the 80 constructs to obtain four expression values corresponding to the four conditions of TF presence/absence – Basal, DL, TWI, DL & TWI.

#### Synthetic constructs for evaluation of distance-dependent interactions

To characterize distance-dependent interactions learned by CoNSEPT models, we examined their predictions on additional synthetic constructs containing pairs of DL-TWI, DL-SNA, TWI-SNA or DL-DL binding sites at progressively increasing separations (from 5bp to 195bp), located at a random location in a 635 bp long sequence. Thus, a collection of 3900 synthetic enhancers was tested, corresponding to 39 different inter-site spacings and 100 different locations of the site pair in the enhancer. The TF binding sites were set to consensus sites of corresponding PWMs and the remaining bases of the enhancers were set to the dummy base “N” (see above). It is worth mentioning that we did not evaluate TWI-TWI distance dependent interactions since the training enhancers contain a maximum of two binding sites for this TF (Figure 1A), potentially preventing the reliable learning of their distance-dependent interactions.

We used the trained CoNSEPT models to predict expression driven by each synthetic construct with relative levels of DL, TWI, and SNA set to of 0.4, 0.3, and 0.4 respectively, reflecting the 6^th^ of the 17 “bins” along the V-D axis. (This bin was selected as it represents the dorsal boundary of SNA expression.) For each TF pair at a specific inter-site spacing, we averaged the predicted expression over all 100 constructs with different placements of the TF pair. We evaluated the GEMSTAT model (NR/COOP) on the same constructs containing pairs of TFs; however, we replaced the dummy bases “N” with random draws from the set of four nucleotides (A, C, G, and T) with equal probability.

### Data and Code Availability

We used the publicly available dataset (including enhancer sequences, gene expressions, TF levels, and PWMs) first published and analyzed by Sayal et al. (14).

The GEMSTAT version allowing “limited contact” scheme used in this study is available at: https://github.com/PayamDiba/Manuscript_tools/tree/main/GEMSTAT_direct_limited.

CoNSEPT is freely available as a python package at: https://github.com/PayamDiba/CoNSEPT.

## Acknowledgement

The authors would like to thank David N. Arnosti and Rupinder Sayal for insightful discussions as well as helping us work with their data.

This work was supported in part by the NIH (grants R01GM114341 and R35GM131819 to SS). Additionally, this work was funded by the DOE Center for Advanced Bioenergy and Bioproducts Innovation (U.S. Department of Energy, Office of Science, Office of Biological and Environmental Research under Award Number DE-SC0018420). Any opinions, findings, and conclusions or recommendations expressed in this publication are those of the author(s) and do not necessarily reflect the views of the U.S. Department of Energy.

## Notes

### Competing Interest Statement

The authors have declared no competing interest.

## References

1. Spitz, F. and Furlong, E.E.M. (2012) Transcription factors: From enhancer binding to developmental control. Nat. Rev. Genet., 13, 613–626.

2. Kazemian, M., Pham, H., Wolfe, S.A., Brodsky, M.H. and Sinha, S. (2013) Widespread evidence of cooperative DNA binding by transcription factors in Drosophila development. Nucleic Acids Res., 41, 8237–8252.

3. Hobert, O. (2008) Gene regulation by transcription factors and MicroRNAs. Science (80-.)., 319, 1785–1786.

4. Hong, J.W., Hendrix, D.A., Papatsenko, D. and Levine, M.S. (2008) How the Dorsal gradient works: Insights from postgenome technologies. Proc. Natl. Acad. Sci. U. S. A., 105, 20072–20076.

5. Jaeger, J., Manu and Reinitz, J. (2012) Drosophila blastoderm patterning. Curr. Opin. Genet. Dev., 22, 533–541.

6. Johnston, D.S. and Nüsslein-Volhard, C. (1992) The origin of pattern and polarity in the Drosophila embryo. Cell, 68, 201–219.

7. Struffi, P., Corado, M., Kulkarni, M. and Arnosti, D.N. (2004) Quantitative contributions of CtBP-dependent and -independent repression activitis of Knirps. Development, 131, 2419–2429.

8. Nibu, Y., Senger, K. and Levine, M. (2003) CtBP-Independent Repression in the Drosophila Embryo. Mol. Cell. Biol., 23, 3990–3999.

9. Nibu, Y. and Levine, M.S. (2001) CtBP-dependent activities of the short-range giant repressor in the Drosophila embryo. Proc. Natl. Acad. Sci. U. S. A., 98, 6204–6208.

10. Bhaskar, V. and Courey, A.J. (2002) The MADF-BESS domain factor Dip3 potentiates synergistic activation by Dorsal and Twist. Gene, 299, 173–184.

11. Szymanski, P. and Levine, M. (1995) Multiple modes of dorsal-bHLH transcriptional synergy in the Drosophila embryo. EMBO J., 14, 2229–2238.

12. King, D.M., Hong, C.K.Y., Shepherdson, J.L., Granas, D.M., Maricque, B.B. and Cohen, B.A. (2020) Synthetic and genomic regulatory elements reveal aspects of Cis-regulatory grammar in mouse embryonic stem cells. Elife, 9.

13. Kulkarni, M.M. and Arnosti, D.N. (2005) cis-Regulatory Logic of Short-Range Transcriptional Repression in Drosophila melanogaster. Mol. Cell. Biol., 25, 3411–3420.

14. Sayal, R., Dresch, J.M., Pushel, I., Taylor, B.R. and Arnosti, D.N. (2016) Quantitative perturbation-based analysis of gene expression predicts enhancer activity in early Drosophila embryo. Elife, 5.

15. White, M.A., Kwasnieski, J.C., Myers, C.A., Shen, S.Q., Corbo, J.C. and Cohen, B.A. (2016) A Simple Grammar Defines Activating and Repressing cis-Regulatory Elements in Photoreceptors. Cell Rep., 17, 1247–1254.

16. Deplancke, B., Alpern, D. and Gardeux, V. (2016) The Genetics of Transcription Factor DNA Binding Variation. Cell, 166, 538–554.

17. Ay, A. and Arnosti, D.N. (2011) Mathematical modeling of gene expression: A guide for the perplexed biologist. Crit. Rev. Biochem. Mol. Biol., 46, 137–151.

18. Fakhouri, W.D., Ay, A., Sayal, R., Dresch, J., Dayringer, E. and Arnosti, D.N. (2010) Deciphering a transcriptional regulatory code: Modeling short-range repression in the Drosophila embryo. Mol. Syst. Biol., 6.

19. Vahrenkamp, J.M., Yang, C.H., Rodriguez, A.C., Almomen, A., Berrett, K.C., Trujillo, A.N., Guillen, K.P., Welm, B.E., Jarboe, E.A., Janat-Amsbury, M.M., et al. (2018) Clinical and Genomic Crosstalk between Glucocorticoid Receptor and Estrogen Receptor α In Endometrial Cancer. Cell Rep., 22, 2995–3005.

20. Farley, E.K., Olson, K.M., Zhang, W., Rokhsar, D.S. and Levine, M.S. (2016) Syntax compensates for poor binding sites to encode tissue specificity of developmental enhancers. Proc. Natl. Acad. Sci. U. S. A., 113, 6508–6513.

21. Janssens, H., Hou, S., Jaeger, J., Kim, A.R., Myasnikova, E., Sharp, D. and Reinitz, J. (2006) Quantitative and predictive model of transcriptional control of the Drosophila melanogaster even skipped gene. Nat. Genet., 38, 1159–1165.

22. Ilsley, G.R., Fisher, J., Apweiler, R., DePace, A.H. and Luscombe, N.M. (2013) Cellular resolution models for even skipped regulation in the entire Drosophila embryo. Elife, 2.

23. Crocker, J., Ilsley, G.R. and Stern, D.L. (2016) Quantitatively predictable control of Drosophila transcriptional enhancers in vivo with engineered transcription factors. Nat. Genet., 48, 292–298.

24. Segal, E., Raveh-Sadka, T., Schroeder, M., Unnerstall, U. and Gaul, U. (2008) Predicting expression patterns from regulatory sequence in Drosophila segmentation. Nature, 451, 535–540.

25. Zinzen, R.P. and Papatsenko, D. (2007) Enhancer Responses to Similarly Distributed Antagonistic Gradients in Development. PLoS Comput. Biol., 3, e84.

26. Reinitz, J., Hou, S. and Sharp, D.H. (2003) Transcriptional Control in Drosophila. Complexus, 1, 54–64.

27. Dresch, J.M., Thompson, M.A., Arnosti, D.N. and Chiu, C. (2013) Two-layer mathematical modeling of gene expression: Incorporating dna-level information and system dynamics. SIAM J. Appl. Math., 73, 804–826.

28. He, X., Samee, M.A.H., Blatti, C. and Sinha, S. (2010) Thermodynamics-based models of transcriptional regulation by enhancers: The roles of synergistic activation, cooperative binding and short-range repression. PLoS Comput. Biol., 6.

29. Samee, M.A.H., Lim, B., Samper, N., Lu, H., Rushlow, C.A., Jiménez, G., Shvartsman, S.Y. and Sinha, S. (2015) A Systematic Ensemble Approach to Thermodynamic Modeling of Gene Expression from Sequence Data. Cell Syst., 1, 396–407.

30. Grah, R., Zoller, B. and Tkačik, G. (2020) Nonequilibrium models of optimal enhancer function. Proc. Natl. Acad. Sci. U. S. A., 117, 31614–31622.

31. Ahsendorf, T., Wong, F., Eils, R. and Gunawardena, J. (2014) A framework for modelling gene regulation which accommodates non-equilibrium mechanisms. BMC Biol., 12, 102.

32. Estrada, J., Wong, F., DePace, A. and Gunawardena, J. (2016) Information Integration and Energy Expenditure in Gene Regulation. Cell, 166, 234–244.

33. Gertz, J., Siggia, E.D. and Cohen, B.A. (2009) Analysis of combinatorial cis-regulation in synthetic and genomic promoters. Nature, 457, 215–218.

34. Beer, M.A. and Tavazoie, S. (2004) Predicting gene expression from sequence. Cell, 117, 185–198.

35. Blatti, C., Kazemian, M., Wolfe, S., Brodsky, M. and Sinha, S. (2015) Integrating motif, DNA accessibility and gene expression data to build regulatory maps in an organism. Nucleic Acids Res., 43, 3998–4012.

36. Avsec, Ž., Weilert, M., Shrikumar, A., Krueger, S., Alexandari, A., Dalal, K., Fropf, R., McAnany, C., Gagneur, J., Kundaje, A., et al. (2021) Base-resolution models of transcription-factor binding reveal soft motif syntax. Nat. Genet., 10.1038/s41588-021-00782-6.

37. Kelley, D.R., Snoek, J. and Rinn, J.L. (2016) Basset: Learning the regulatory code of the accessible genome with deep convolutional neural networks. Genome Res., 26, 990–999.

38. Alipanahi, B., Delong, A., Weirauch, M.T. and Frey, B.J. (2015) Predicting the sequence specificities of DNA- and RNA-binding proteins by deep learning. Nat. Biotechnol., 33, 831–838.

39. Zhou, J. and Troyanskaya, O.G. (2015) Predicting effects of noncoding variants with deep learning-based sequence model. Nat. Methods, 12, 931–934.

40. Kazemian, M., Blatti, C., Richards, A., Mccutchan, M. and Wakabayashi-Ito, N. (2010) Quantitative Analysis of the Drosophila Segmentation Regulatory Network Using Pattern Generating Potentials. PLoS Biol, 8, 1000456.

41. Samee, M.A.H., Lydiard-Martin, T., Biette, K.M., Vincent, B.J., Bragdon, M.D., Eckenrode, K.B., Wunderlich, Z., Estrada, J., Sinha, S. and DePace, A.H. (2017) Quantitative Measurement and Thermodynamic Modeling of Fused Enhancers Support a Two-Tiered Mechanism for Interpreting Regulatory DNA. Cell Rep., 21, 236–245.

42. Shirokawa, J.M. and Courey, A.J. (1997) A direct contact between the dorsal rel homology domain and Twist may mediate transcriptional synergy. Mol. Cell. Biol., 17, 3345–3355.

43. Nibu, Y., Zhang, H. and Levine, M. (1998) Interaction of short-range repressors with Drosophila CtBP in the embryo. Science (80–.)., 280, 101–104.

44. Nibu, Y., Zhang, H., Bajor, E., Barolo, S., Small, S. and Levine, M. (1998) dCtBP mediates transcriptional repression by Knirps, Kruppel and Snail in the Drosophila embryo. EMBO J., 17, 7009–7020.

45. Chinnadurai, G. (2002) CtBP, an unconventional transcriptional corepressor in development and oncogenesis. Mol. Cell, 9, 213–224.

46. Struffi, P. and Arnosti, D.N. (2005) Functional interaction between the Drosophila knirps short range transcriptional repressor and RPD3 histone deacetylase. J. Biol. Chem., 280, 40757–40765.

47. Swanson, C.I., Evans, N.C. and Barolo, S. (2010) Structural rules and complex regulatory circuitry constrain expression of a Notch- and EGFR-regulated eye enhancer. Dev. Cell, 18, 359–370.

48. Crocker, J., Tamori, Y. and Erives, A. (2008) Evolution Acts on Enhancer Organization to Fine-Tune Gradient Threshold Readouts. PLoS Biol., 6, e263.

49. Arnold, C.D., Gerlach, D., Stelzer, C., Boryń, Ł.M., Rath, M. and Stark, A. (2013) Genome-wide quantitative enhancer activity maps identified by STARR-seq. Science (80–.), 339, 1074–1077.

50. Maricque, B.B., Dougherty, J.D. and Cohen, B.A. (2017) A genome-integrated massively parallel reporter assay reveals DNA sequence determinants of cis-regulatory activity in neural cells. Nucleic Acids Res., 45.

51. Melnikov, A., Zhang, X., Rogov, P., Wang, L. and Mikkelsen, T.S. (2014) Massively parallel reporter assays in cultured mammalian cells. J. Vis. Exp., 10.3791/51719.

52. Gertz, J., Siggia, E.D. and Cohen, B.A. (2009) Analysis of combinatorial cis-regulation in synthetic and genomic promoters. Nature, 457, 215–218.

53. Papatsenko, D. and Levine, M.S. (2008) Dual regulation by the Hunchback gradient in the Drosophila embryo. Proc. Natl. Acad. Sci. U. S. A., 105, 2901–2906.

54. Kim, A.R., Martinez, C., Ionides, J., Ramos, A.F., Ludwig, M.Z., Ogawa, N., Sharp, D.H. and Reinitz, J. (2013) Rearrangements of 2.5 Kilobases of Noncoding DNA from the Drosophila even-skipped Locus Define Predictive Rules of Genomic cis-Regulatory Logic. PLoS Genet., 9.

55. Gray, I., Szymanski, P. and Levine, M. (1994) Short-range repression permits multiple enhancers to function autonomously within a complex promoter. Genes Dev., 8, 1829– 1838.

56. Courey, A.J. and Jia, S. (2001) Transcriptional repression: The long and the short of it. Genes Dev., 15, 2786–2796.

57. Settles, B. (2012) Active learning. Synth. Lect. Artif. Intell. Mach. Learn., 18.

58. Khajouei, F. and Sinha, S. (2018) An information theoretic treatment of sequence-to-expression modeling. PLOS Comput. Biol., 14, e1006459.

59. Lal, A., Chiang, Z.D., Yakovenko, N., Duarte, F.M., Israeli, J. and Buenrostro, J.D. (2019) AtacWorks: A deep convolutional neural network toolkit for epigenomics. bioRxiv, 10.1101/829481.

60. Angermueller, C., Lee, H.J., Reik, W. and Stegle, O. (2017) DeepCpG: Accurate prediction of single-cell DNA methylation states using deep learning. Genome Biol., 18, 67.

61. Agarwal, V. and Shendure, J. (2020) Predicting mRNA Abundance Directly from Genomic Sequence Using Deep Convolutional Neural Networks. Cell Rep., 31, 107663.

62. Weirauch, M.T., Yang, A., Albu, M., Cote, A.G., Montenegro-Montero, A., Drewe, P., Najafabadi, H.S., Lambert, S.A., Mann, I., Cook, K., et al. (2014) Determination and inference of eukaryotic transcription factor sequence specificity. Cell, 158, 1431–1443.

63. Liu, Y., Barr, K. and Reinitz, J. (2020) Fully interpretable deep learning model of transcriptional control. Bioinformatics, 36, i499–i507.

64. Khajouei, F., Samper, N., Djabrayan, N.J., Lunt, B., Jiménez, G. and Sinha, S. (2020) Model-based analysis of polymorphisms in an enhancer reveals cis-regulatory mechanisms. bioRxiv, 10.1101/2020.02.07.939264.

65. Tabe-Bordbar, S., Song, Y.J., Lunt, B.J., Prasanth, K. V. and Sinha, S. (2020) Mechanistic analysis of enhancer sequences in the Estrogen Receptor transcriptional program. bioRxiv, 10.1101/2020.11.08.373555.

66. Duque, T., Samee, M.A.H., Kazemian, M., Pham, H.N., Brodsky, M.H. and Sinha, S. (2014) Simulations of Enhancer Evolution Provide Mechanistic Insights into Gene Regulation. Mol. Biol. Evol., 31, 184–200.

67. Srivastava, N., Hinton, G., Krizhevsky, A. and Salakhutdinov, R. (2014) Dropout: A Simple Way to Prevent Neural Networks from Overfitting.

68. Ba, J.L., Kiros, J.R. and Hinton, G.E. (2016) Layer Normalization. arXiv Prepr.

69. Kingma, D.P. and Ba, J.L. (2014) Adam: A method for stochastic optimization. In arXiv preprint. International Conference on Learning Representations, ICLR.

